# Stepwise origin and evolution of a cryptic antimicrobial peptide in mammalian lactoferrin

**DOI:** 10.64898/2026.02.05.703623

**Authors:** Titas Sil, Caitlin H. Kowalski, Sierra Scamfer, Matthew F. Barber

**Affiliations:** Institute of Ecology and Evolution, University of Oregon; Eugene, OR 97403, USA; Department of Biology, University of Oregon; Eugene, OR 97403, USA; Department of Dermatology, Dartmouth Hitchcock Medical Center; Lebanon, NH 03756, USA; Institute of Molecular Biology, University of Oregon; Eugene, OR 97403, USA

**Author notes:** **Author Contributions:** T.S. and M.F.B. designed research; T.S. performed research; T.S., C.H.K., and S.S. contributed to data collection; T.S. and M.F.B. wrote the paper; T.S., C.H.K., S.S., and M.F.B edited the paper. **Competing Interest Statement:** Authors declare that they have no competing interests.

**Keywords:** protein evolution, antimicrobial peptides, innate immunity, pathogens, lactoferrin

## Abstract

Antimicrobial peptides (AMPs) constitute key components of innate immunity across the tree of life. Canonical AMPs are typically translated as small proteins and secreted from host cells to act against microbes. However, cryptic AMP-like domains are also embedded within diverse proteins not classically associated with antimicrobial function. How such embedded AMPs first emerge and diversify remains unclear. Here we retrace the origin and evolution of the abundant mammalian protein lactoferrin and its embedded AMP, lactoferricin. By resurrecting extinct lactoferrin ancestors dating back to the earliest mammals, we identify a gradual enrichment of cationic and hydrophobic amino acids in the lactoferricin domain over time. These changes enabled ancient lactoferricin to first rupture bacterial membranes, an activity that was later enhanced in extant mammals conferring potent bactericidal activity against diverse bacteria. In addition, we find that recent natural selection within the lactoferricin domain has continued to modulate antimicrobial activity on recent evolutionary timescales. In particular, we pinpoint a single rapidly evolving site in lactoferricin among great apes that significantly enhances antimicrobial potency against major pathogenic bacteria. Together our study illustrates how novel immune protein functions can arise, evolve, and diversify to strengthen host defense against diverse microbial pathogens.

## Introduction

The evolution of immune function is vital for species survival in response to pathogen antagonism ^1–3^. Antimicrobial peptides (AMPs), also known as host defense peptides, are major contributors to innate immunity across all domains of life, displaying a range of antimicrobial and immunomodulatory activities ^4–7^. Well-characterized AMPs are typically small (less than 50 amino acids) and positively charged, promoting interactions with negatively charged microbial cell envelopes ^4,5,8^. These interactions can lead to membrane disruption, cell lysis, and impairment of essential cellular processes, ultimately inhibiting microbial growth or triggering cell death ^8^. Beyond canonical AMPs, an increasing number of studies have identified the presence of AMP-like domains embedded within larger proteins ^9–13^. These domains can in some cases be liberated through proteolytic cleavage, suggesting a mechanism by which diverse proteins may acquire immune functionality. At present, little is known regarding the origins of these embedded AMP domains or the molecular changes that enabled the acquisition of antimicrobial activity.

The mammalian protein lactoferrin provides an exemplar for the evolution of antimicrobial functions. Lactoferrin is an abundant secreted iron-binding protein that arose by duplication of the transferrin gene in the ancestor of placental mammals approximately 160 million years ago (Fig. S1A) ^14,15^. Transferrin proteins are typically composed of two homologous domains, termed the N- and C-lobes, each of which binds a single ferric iron ion with high affinity ^15–18^. Serum transferrin mediates iron transport in the bloodstream, delivering this essential metal nutrient to cells via receptor-mediated endocytosis ^15^. Transferrin family proteins can also provide a defensive benefit by sequestering iron from invasive pathogens, a process termed “nutritional immunity”^19–21^. Lactoferrin, in contrast to transferrin, is abundantly expressed in mammalian milk, colostrum, tears, secondary granules of neutrophils, and other body fluids ^22,23^. Since its birth and divergence, lactoferrin has also acquired novel antimicrobial functions which are absent in extant transferrin (Fig. S1A-B) ^23^. The N-terminal region of the lactoferrin N-lobe can be proteolytically cleaved by abundant host proteases such as pepsin, trypsin, and chymotrypsin to generate cationic AMPs, including lactoferricin and lactoferrampin ^24–29^. These peptides are capable of killing diverse bacterial and fungal pathogens as well as inhibiting microbial biofilm formation ^26,29– 32^. Lactoferrin-derived AMPs share key features with conventional AMPs, including a high proportion of positively charged and hydrophobic amino acids that confer amphipathic properties (Fig. 1A, S2A). Despite its functional importance, the evolutionary origin of antimicrobial activity within the lactoferricin domain remains poorly understood. Previously we found that lactoferricin exhibits evidence of repeated positive selection in simian primates, with codons possessing an elevated rate of non-synonymous to synonymous substitutions (dN/dS) ^33^. This suggests that lactoferricin has continued to diversify within mammals since the acquisition of antimicrobial activity. Determining how this novel function arose and subsequently evolved in an otherwise well-conserved iron-transport protein could offer broader insights into the origin of antimicrobial proteins and the mechanisms by which new biological functions arise. Here we retrace the stepwise emergence of lactoferricin antimicrobial activity in ancestral mammals, as well as demonstrate how recent natural selection has enhanced the function of this abundant host defense protein.

**Figure 1.**
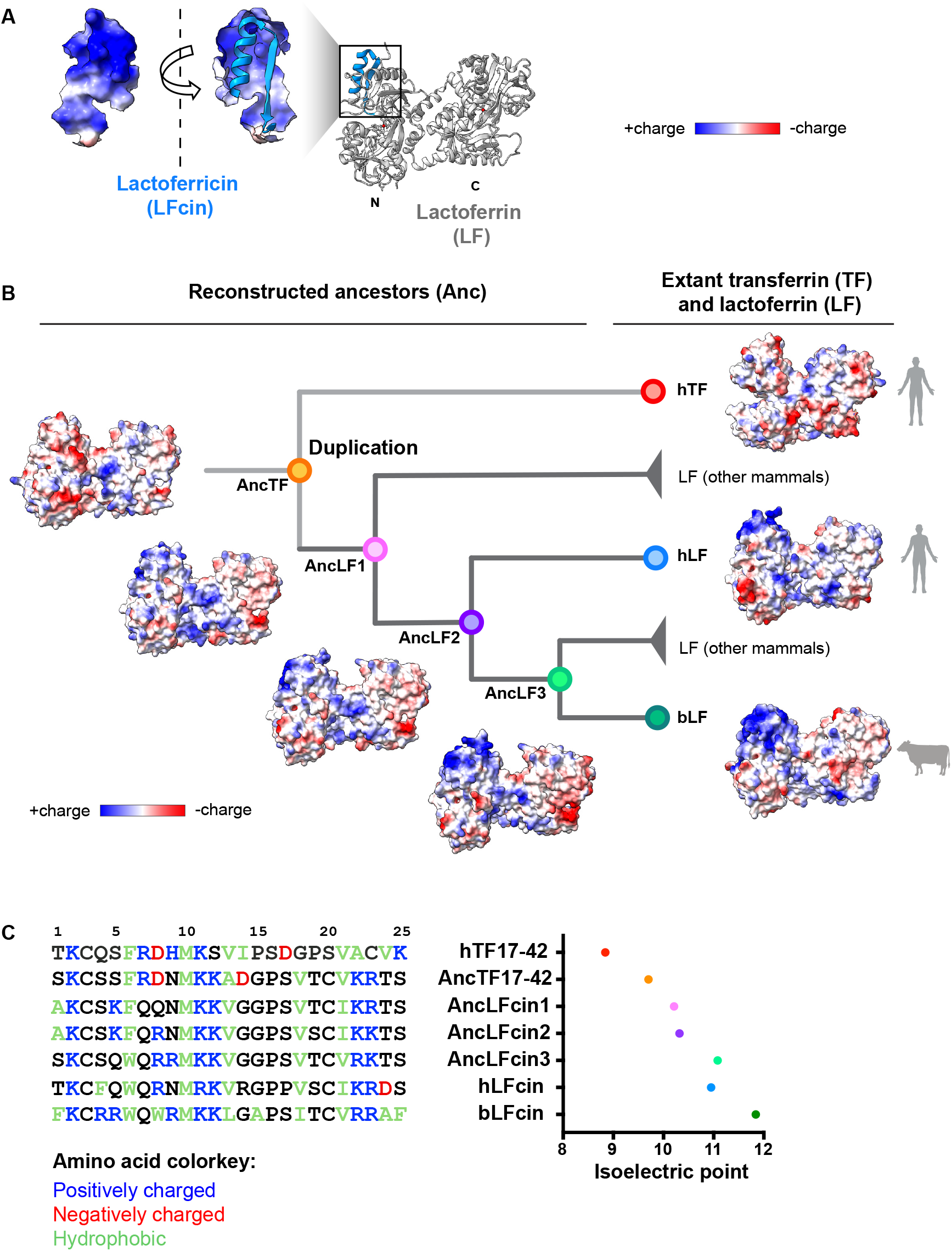
Retracing the emergence of antimicrobial function in mammalian lactoferrin. (**A**) Structure of the N-terminal lactoferricin region of lactoferrin (PDB: 1lfg), enriched with positively charged amino acids. (**B**) Phylogenetic tree of reconstructed ancestral lactoferrin homologs (AncTF and AncLF) with extant human transferrin (hTF, PDB: 3qyt), human lactoferrin (hLF, PDB: 1lfg), and bovine lactoferrin (bLF, PDB: 1blf) illustrated with surface electrostatic potentials. Structures of ancestral proteins were predicted using AlphaFold2. AncTF represents the last common ancestor of mammalian transferrin and lactoferrin, after which the first lactoferrin ancestor (AncLF1) arose by gene duplication. AncLF2 is the common ancestor of hLF and bLF, while AncLF3 represents a bLF ancestor. (**C**) Alignment of the lactoferricin region from hTF, AncTF, selected AncLFs, hLF, and bLF, with their corresponding isoelectric points. Higher isoelectric points indicate higher incidence of positive charges.

## Results

### Ancestral lactoferrin accumulated positive charges after gene duplication

To investigate the origin and evolutionary history of the lactoferricin domain, we first compared lactoferricin sequences across diverse mammals (Fig. S2A). Given that lactoferricin peptides have been reported of different lengths in different species, we focused our studies on the core 25 amino acids conserved across lactoferricin orthologs (amino acids 17-42 in mature human lactoferrin) ^24,25^. Most homologs possessed elevated isoelectric points indicating enrichment of positively charged amino acids such as arginine (R), lysine (K), and histidine (H). They also contained interspersed hydrophobic residues characteristic of AMPs. Despite these conserved physiochemical properties, lactoferricin amino acid sequences were highly diverse across orthologs, making it difficult to pinpoint amino acids that may have been responsible for antimicrobial function (Fig. S2A).

To retrace the mutations that led to the emergence of a novel antimicrobial function, we reconstructed ancestral lactoferrin and transferrin sequences across diverse mammals (Fig. 1B). Ancestral sequence reconstruction provides a powerful approach to characterize the function of ancient proteins ^34–36^. Using the Topiary software pipeline ^37^ and publicly available transferrin and mammalian lactoferrin sequences from NCBI, we predicted ancestral amino acid sequences of transferrin and lactoferrin proteins (Fig. S2B). We focused on ancestral proteins (prefixed “Anc”) at key nodes of the phylogenetic tree where the lactoferricin domain underwent significant divergence (Fig. 1B-C). Each predicted sequence was associated with posterior probability scores, which identified the most likely residue at each site as well as the most probable alternative (Fig. S3). These data were used to generate corresponding alternative (Altall_Anc) peptide sequences, in which alternative amino acid states with a posterior probability greater than 0.2 replaced the most probable ancestral amino acid. To assess the robustness of our findings, we included both the primary ancestral reconstructions and their alternate versions in subsequent functional analyses (Fig. S4).

To compare the antimicrobial function of lactoferricin between ancestral and extant variants, we chose human and bovine lactoferrin (hLF and bLF, respectively) as extant references, given the extensive previous work on these orthologs in the field ^22,24,25,27^. As a negative control we selected human serum transferrin (hTF) as it did not exhibit antimicrobial function in the N-terminal region (Fig. S1B). We referred to the last common ancestor of mammalian transferrin prior to the gene duplication event as AncTF. Gene duplication produced the first ancestral lactoferrin, which we designated AncLF1 (Fig. 1B). We also considered two intermediate ancestors, denoted AncLF2 and AncLF3. AncLF2 represents the last common ancestor of hLF and bLF in the gene phylogeny, and AncLF3 represents a bovine-specific ancestor that accumulated multiple substitutions in the lactoferricin region relative to AncLF2 (Fig. 1C). We then predicted the structures of these full-length ancestral proteins using AlphaFold2 (Fig. 1B). When the surface electrostatic potentials were compared, we observed a progressive increase in positive charge density (blue surface) in the N-terminal region of lactoferrin following the duplication event (Fig. 1B). This suggests that the emergence of positively charged surfaces in the N-terminus occurred shortly before or after the origin of lactoferrin itself in placental mammals.

We next examined the lactoferricin domains of these reconstructed lactoferrin ancestors, which we refer to as AncLFcin1, AncLFcin2, and AncLFcin3 (Fig. 1C). As the corresponding region in transferrin (residues 17-42) does not form a known antimicrobial peptide, we referred to these homologous sequences as AncTF17-42 in ancestral transferrin and hTF17-42 in human serum transferrin. We observed that cationic amino acids became more abundant in lactoferricin post-duplication, with the emergence of a K5 substitution in AncLFcin1, R8 in AncLFcin2, and R9 in AncLFcin3 (Fig. 1C). To quantify this trend, we calculated peptide isoelectric points, which increased from AncTF17-42 to bLFcin (Fig. 1C). Furthermore, we noted the emergence of hydrophobic residues adjacent to positively charged amino acids in the lactoferricin domain after duplication (e.g., alanine 1 in AncLFcin1 and AncLFcin2). Based on these observations, we hypothesized that the increase in positive charge and hydrophobicity enabled lactoferricin to acquire antimicrobial function by mediating association with bacterial cell envelopes.

### Early origin of antimicrobial activity in lactoferricin

To characterize the antimicrobial potency of ancestral and extant lactoferricin domains, we synthesized these peptides and tested their activity against several bacterial pathogens (Fig. 2). Among Gram-negative bacteria we included *Pseudomonas aeruginosa*, a pathogen frequently associated with nosocomial infections, chronic wounds, and cystic fibrosis, as well as a reference strain of the enteric bacterium *Escherichia coli* ^38–40^. We also tested representative Gram-positive bacteria including *Staphylococcus aureus*, a major cause of invasive hospital and community-acquired infections, as well as *Streptococcus agalactiae*, a frequent agent of severe infections in neonates ^41–43^.

**Figure 2.**
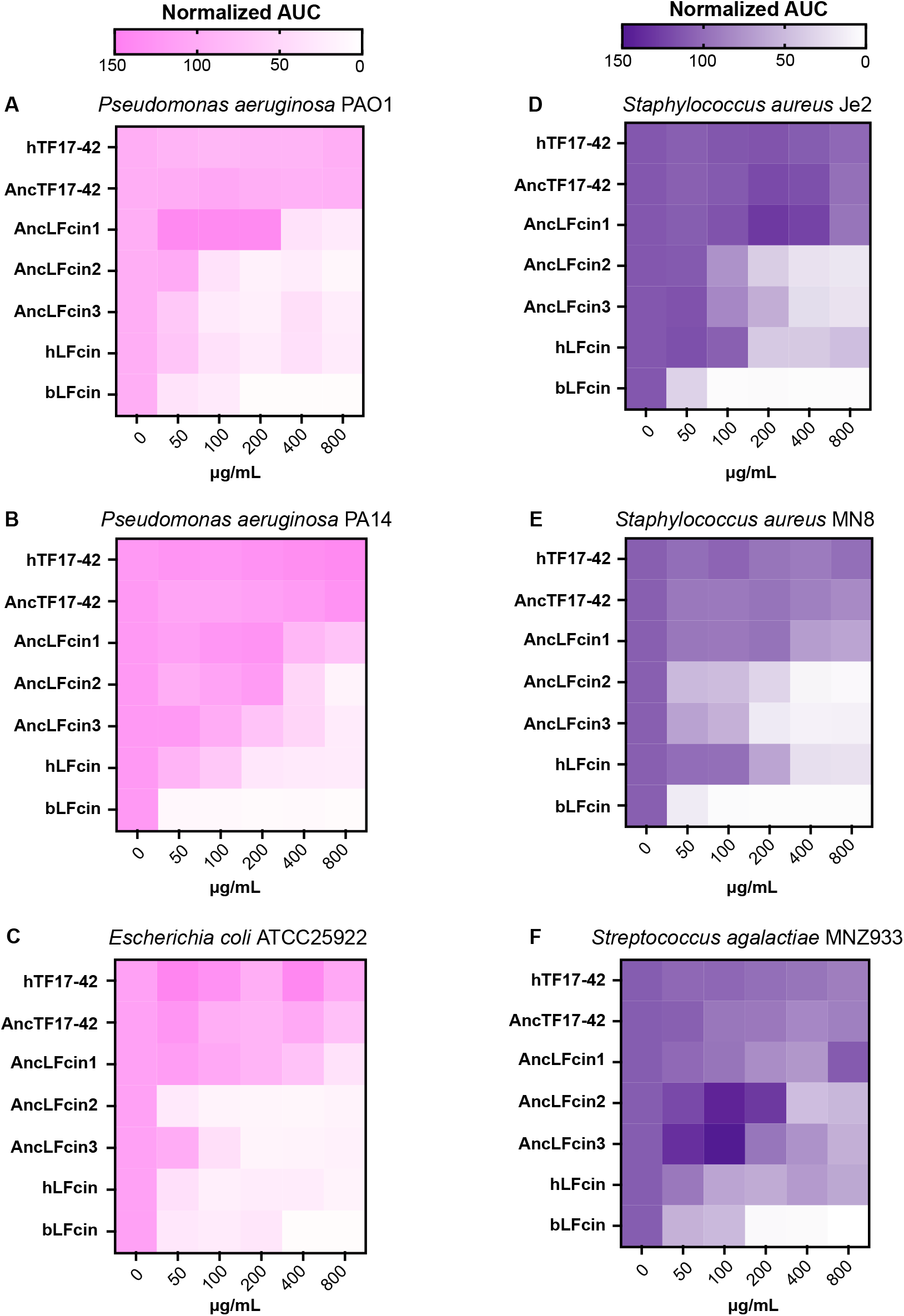
origin of antimicrobial activity in lactoferricin. Heatmaps depict area under the curve (AUC) values normalized against the untreated control from bacterial growth curves performed in the presence of indicated peptides at specified concentrations. Gram-negative strains are shown in pink (A-C), Gram-positive strains in purple (D-F).

To compare the antimicrobial potency of ancestral and extant lactoferricin homologs, we measured bacterial growth in presence of the peptides at various concentrations over 24 hours. Human lactoferrin concentrations range widely *in vivo* from 7-8 mg/mL in colostrum, 1-3 mg/mL in breast milk, 0.1-2 mg/mL in tears and saliva, and even lower levels (0.001-0.1 mg/mL) in other secretory body fluids ^44^. We therefore used physiologically relevant concentrations of each peptide, ranging from 0.1-1mg/mL, to assess their dose-dependent antimicrobial activities. Bacterial growth at each concentration was measured by normalized area under the curve (AUC) calculations, where higher AUC corresponds to increased bacterial growth. Relative to the untreated control, peptides exhibiting <50% bacterial growth were classified as having high antimicrobial potency, those with 50–75% growth as moderate, and those with >75% growth as weak. We observed that all bacteria grew to high levels in the presence of hTF17-42, indicating it has little to no antimicrobial activity as expected (Fig. 2). Similarly, the ancestral-derived peptide AncTF17-42 exhibited negligible antimicrobial activity across all doses and pathogens (Fig. 2). AncLFcin1, in contrast, reduced growth by >50% at higher doses against a subset of pathogens, for example at 400 µg/mL against *P. aeruginosa* PAO1 and at 800 µg/mL against *E. coli* (Fig. 2A-B). These findings suggest that antimicrobial activity in this domain arose after the gene duplication event that gave rise to mammalian lactoferrin.

AncLFcin2, representing the last common ancestor of hLF and bLF, exhibited higher potency at lower concentrations across all pathogens tested relative to AncLFcin1 (Fig. 2). For example, it was highly potent at 100 µg/mL against *P. aeruginosa* PAO1, and at 50 µg/mL against *E. coli* and *S. aureus* MN8 (Fig. 2A, 2C, 2E). Against other pathogens, AncLFcin2 also reduced bacterial growth by >50% in the 100-400 µg/mL range. These results suggest that antimicrobial activity of the lactoferricin domain was substantially enhanced between AncLFcin1 and AncLFcin2. AncLFcin3 exhibited broad antimicrobial activity in the 100-800 µg/mL range but overall was less effective than bLFcin. To validate the robustness of our ancestral sequence reconstruction, we also tested alternate versions of each ancestor against *P. aeruginosa* and *S. aureus* (Fig. S4). Overall trends in antimicrobial activity were largely consistent with those observed for the primary reconstructed ancestors. The only exception was Altall_AncTF17-42, which exhibited weak activity against *S. aureus* at the highest concentrations (Fig. S4).

Among extant lactoferricin orthologs, bLFcin was consistently more potent than hLFcin against all pathogens tested, significantly reducing bacterial growth at concentrations as low as 50 µg/mL (Fig. 2). Notably, hLFcin was more effective against Gram-negative bacteria where it restricted growth at 100-200 µg/mL. Against Gram-positive bacteria, the antimicrobial potency of hLFcin only became evident at concentrations above 200 µg/mL. Interestingly, AncLFcin2 was more effective than hLFcin against Gram-positive bacteria. Overall these data indicate that modest antimicrobial activity first emerged in AncLFcin1, immediately after the emergence of lactoferrin in ancient mammals, which then increased in subsequent ancestors. However, given that hLFcin exhibited weaker antimicrobial activity than some earlier ancestors, our findings also suggest that the evolution of lactoferricin antimicrobial function did not increase uniformly during mammalian evolution.

### Stepwise enhancement of bactericidal activity during lactoferricin evolution

After broadly characterizing the antimicrobial nature of the ancestral and extant lactoferricins, we assessed their function in greater detail by measuring growth kinetics and viability of representative Gram-negative (*P. aeruginosa*) and Gram-positive (*S. aureus*) bacteria in the presence of the peptides (Fig. 3). In addition to analysing growth curves and calculating AUCs, we quantified bacterial survival by measuring colony-forming units (CFUs) over time. Based on the differences in dose-dependent activity observed previously, we used 100 µg/mL of the peptides against *P. aeruginosa* and 800 µg/mL against *S. aureus* in subsequent experiments.

**Figure 3.**
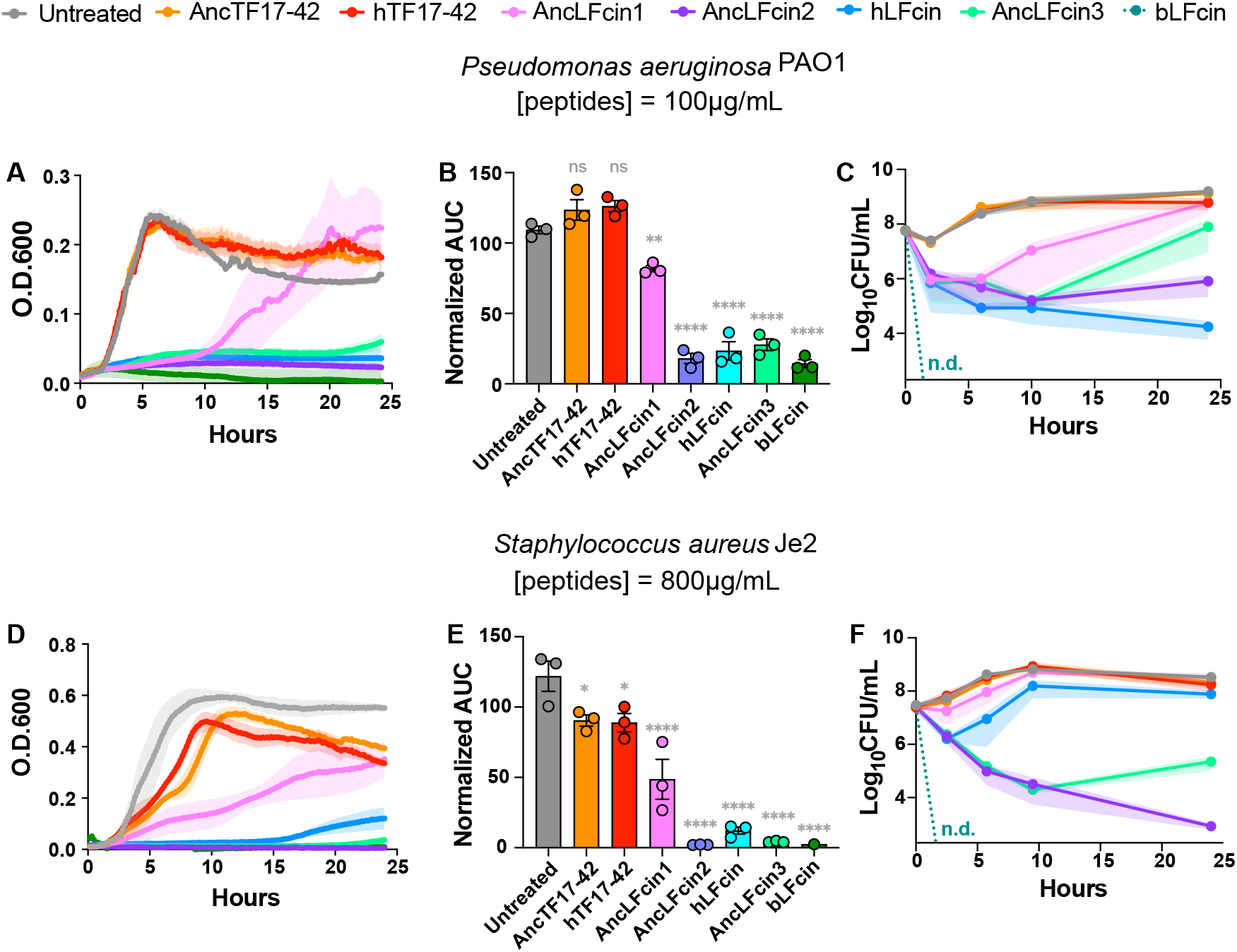
Stepwise evolution of bactericidal activity in ancestral lactoferricin. (**A-C**) *Pseudomonas aeruginosa* strain PAO1 treated with 100 μg/mL of indicated peptides, (**D-F**) *Staphylococcus aureus* strain JE2 treated with 800 μg/mL of indicated peptides. (**A, D**) Growth curves measured by optical density at 600 nm over time. (**B, E**) Corresponding area under the curve (AUC) measurements normalized against untreated control based on bacterial growth curves. (**C, F**) Bacterial survival curves measured as colony forming units (CFUs). All data represent mean and standard error based on three biological replicates. Statistical significance was determined using one-way ANOVA (\**P* ≤ 0.05, \*\**P* ≤ 0.01, \*\*\**P* ≤ 0.001, \*\*\*\**P* ≤ 0.0001; ns, not significant).

Consistent with our previous results, extant and ancestral transferrin-derived peptides, hTF17-42 and AncTF17-42 respectively, exhibited no detectable effects on *P. aeruginosa* growth and survival (Fig. 3A–C). AncLFcin1 displayed an intermediate phenotype, with a modest but significant reduction in overall *P. aeruginosa* growth (Fig. 3A, B). We also observed a rapid decrease in *P. aeruginosa* viability within 2 hours of treatment, followed by a partial recovery (Fig. 3C). In contrast, AncLFcin2 consistently suppressed *P. aeruginosa* growth throughout the experiment, resulting in a significant 1000-fold reduction in bacterial viability after 24 hours of treatment (Fig. 3A-C). AncLFcin3, the ancestor of bLFcin, inhibited *P. aeruginosa* growth and caused 100-1000 fold reduction in viability during the first 15 hours of treatment, after which bacterial growth resumed. hLFcin also consistently suppressed *P. aeruginosa* growth, causing a 10,000 fold reduction in viability at the end of the treatment. Finally, bLFcin exhibited the most potent bactericidal effect, reflected by minimal AUC values and no detectable *P. aeruginosa* colonies within 2 hours of treatment (Fig. 3A-C).

Lactoferricin activity profiles against *S. aureus* differed from those of *P. aeruginosa* in several ways (Fig. 3D–F). hTF17-42 and AncTF17-42 exhibited modest inhibitory effects on *S. aureus* later in the treatment period (Fig. 3D, E), but these differences did not translate into reductions in bacterial viability as measured by CFU counts (Fig. 3F). AncLFcin1 exerted partial inhibition against *S. aureus*, with little reduction in bacterial viability. AncLFcin2 strongly suppressed bacterial growth, resulting in an approximately 100,000 fold reduction in *S. aureus* viability by the end of treatment. AncLFcin3 also significantly inhibited *S. aureus* growth and reduced its viability over the course of the experiment. In contrast, hLFcin initially inhibited bacterial growth and caused a rapid 100-fold reduction in viability. Subsequently, *S. aureus* resumed growth, although this was not reflected in the optical density, possibly due to a reduction in cell size. Again, bLFcin remained the most potent peptide, eliminating detectable colonies within two hours (Fig. 3F). Collectively these findings indicate that weak antimicrobial activity emerged in the earliest lactoferricin ancestor, with bactericidal activity substantially enhanced in subsequent orthologs.

### Lactoferricin membrane permeabilizing activity arose early after gene birth

While both ancestral and extant lactoferricin peptides exhibited some degree of bactericidal activity, they varied dramatically in their potency (Fig. 3). This led us to further examine the molecular mechanisms underlying differences in their activities. A common feature of many cationic AMPs is their ability to disrupt and permeabilize bacterial membranes ^4,6^. To investigate this function, we measured the membrane permeabilizing ability of ancestral and extant lactoferricin peptides against *P. aeruginosa*. Membrane permeabilization was measured by staining with propidium iodide (PI) which enters the bacterial cell and binds DNA only when cell membranes are compromised ^45^. We found that all ancestral and extant lactoferricins, but not transferrin-derived peptides, were capable of permeabilizing bacterial membranes within 30 minutes of incubation as detected by PI staining (Fig. 4A). This finding indicates that bacterial membrane permeabilization was an early trait that emerged in the lactoferricin domain shortly after gene duplication.

**Figure 4.**
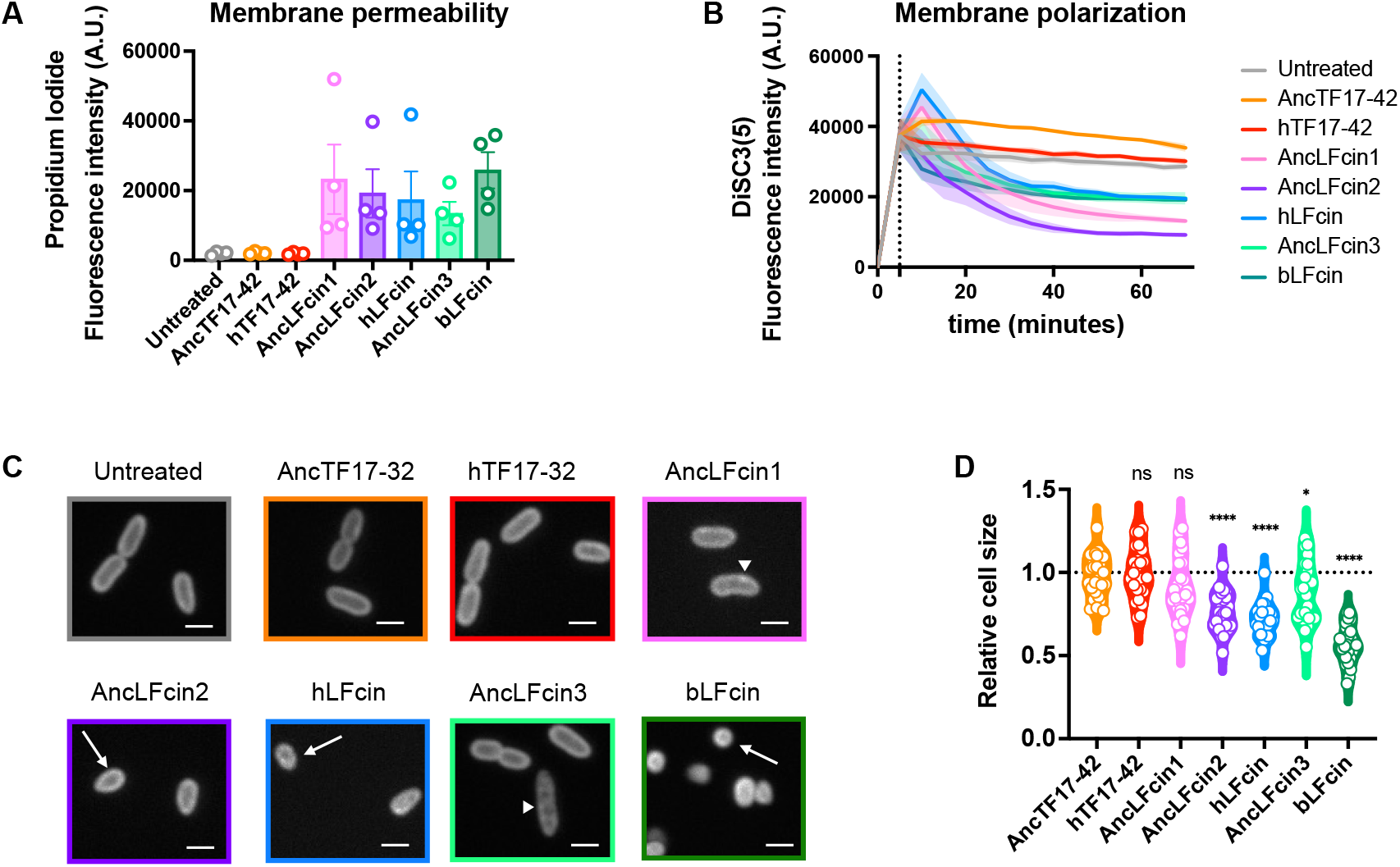
Lactoferricin membrane permeabilizing activity emerged early after gene duplication. *Pseudomonas aeruginosa* strain PAO1 was treated with 100 μg/mL of indicated peptides for 30-60 minutes to assess membrane disruption and cell morphology. (**A**) Membrane permeabilization measured by propidium iodide staining after 30 minutes of treatment. (**B**) Membrane polarization measured by DiSC3(5) dye after 1 hour of treatment. Peptides were added 5 minutes after addition of the dye (shown with the dotted line). Lower fluorescence indicates hyperpolarization. Data represent mean and standard error based on three biological replicates. (**C**) Confocal microscopy images of peptide-treated cells at 30 minutes (scale bar, 1μM). (**D**) Quantification of bacterial cell size from microscopy images (n=100). Statistical significance was determined using one-way ANOVA (\**P* ≤ 0.05, \*\**P* ≤ 0.01, \*\*\**P* ≤ 0.001, \*\*\*\**P* ≤ 0.0001; ns, not significant).

Membrane permeabilization by AMPs can induce bacterial stress responses as well as alter membrane polarization and ion flux, which in turn impacts cell physiology and morphology ^46,47^. To determine the extent to which ancestral and extant lactoferricins affect target cell physiology, we measured bacterial membrane polarization. *P. aeruginosa* cells were treated with the cationic dye 3,3’-dipropylthiadicarbocyanine iodide (DiSC_3_(5)) that accumulates in polarized membranes quenching its fluorescence ^48^. Upon membrane depolarization, DiSC_3_(5) is released, and fluorescence increases. We found that all ancestral and extant lactoferricin-treated cells were hyperpolarized, as demonstrated by a reduction in fluorescence, compared to untreated, hTF17-42, or AncTF17-42-treated cells (Fig. 4B).

Given that all lactoferricin peptides exhibited both membrane-permeabilizing and polarization-altering activities, we sought to understand how these peptides nonetheless differed in their overall antimicrobial potency. To address this, we shifted from population-level assays to single-cell morphological characterization. We hypothesized that lactoferricin variants with increased antimicrobial activity would exert greater impact on bacterial cell envelope integrity. To visualize this, we treated *P. aeruginosa* with the lipophilic dye Nile red, which homogenously binds to membranes and produces brightly stained foci in regions of increased fluidity, indicative of membrane perturbation^45^. Using spinning-disc confocal microscopy, hTF17-42 and AncTF17-42 exhibited no visible effects on bacterial morphology (Fig. 4C). In contrast, cells treated with AncLFcin1, AncLFcin2, or AncLFcin3 displayed puncta-like structures on their membranes, suggesting localized membrane disturbance. These cells also appeared significantly smaller than untreated controls. This effect was most pronounced with hLFcin and bLFcin, where cells became markedly reduced in size, rounded, and membranes were barely distinguishable from the cytosol (Fig. 4C). Quantitative measurements of cell size confirmed that ancestral lactoferricins induced moderate but significant cell shrinkage, whereas extant variants amplified this effect (Fig. 4D).

Membrane disruption is a common mechanism of AMP function, which can cause leaking of cytoplasmic content and shrinkage of cells ^45,49,50^. Such perturbations can be transient and resealable or irreversible, depending on the extent of the damage ^50,51^. Our results suggest that bacteria are able to recover from ancestral lactoferricin-induced membrane damage, whereas extant variants, particularly bLFcin, caused irreversible cell envelope collapse. Together, these findings indicate that the lactoferricin domain initially possessed membrane-permeabilizing activity, which intensified during evolution to produce potent bactericidal effects.

### Rapid evolution of lactoferricin antimicrobial activity in primates

In addition to investigating the molecular origins of the lactoferricin domain, we also sought to determine how lactoferricin antimicrobial activity has changed over recent evolutionary timescales. Previous work by our group found evidence that lactoferrin has been subject to repeated positive selection within human populations as well as between simian primate species, suggesting antimicrobial activity may have been modulated via recurrent adaptation ^18,33^. We identified several sites in the N-lobe of lactoferrin with elevated rates of nonsynonymous to synonymous substitutions (dN/dS), including two within the lactoferricin region (Fig. 5A, S5) ^33^. For example, position 5 varies between primates, with humans and many monkeys encoding a glutamine (Q), whereas all other great apes encode an arginine (R). In addition, position 12 is variable among primates as well as polymorphic within human populations, generally toggling between cationic arginine and lysine (K12 or R12) residues ^33^.

**Figure 5.**
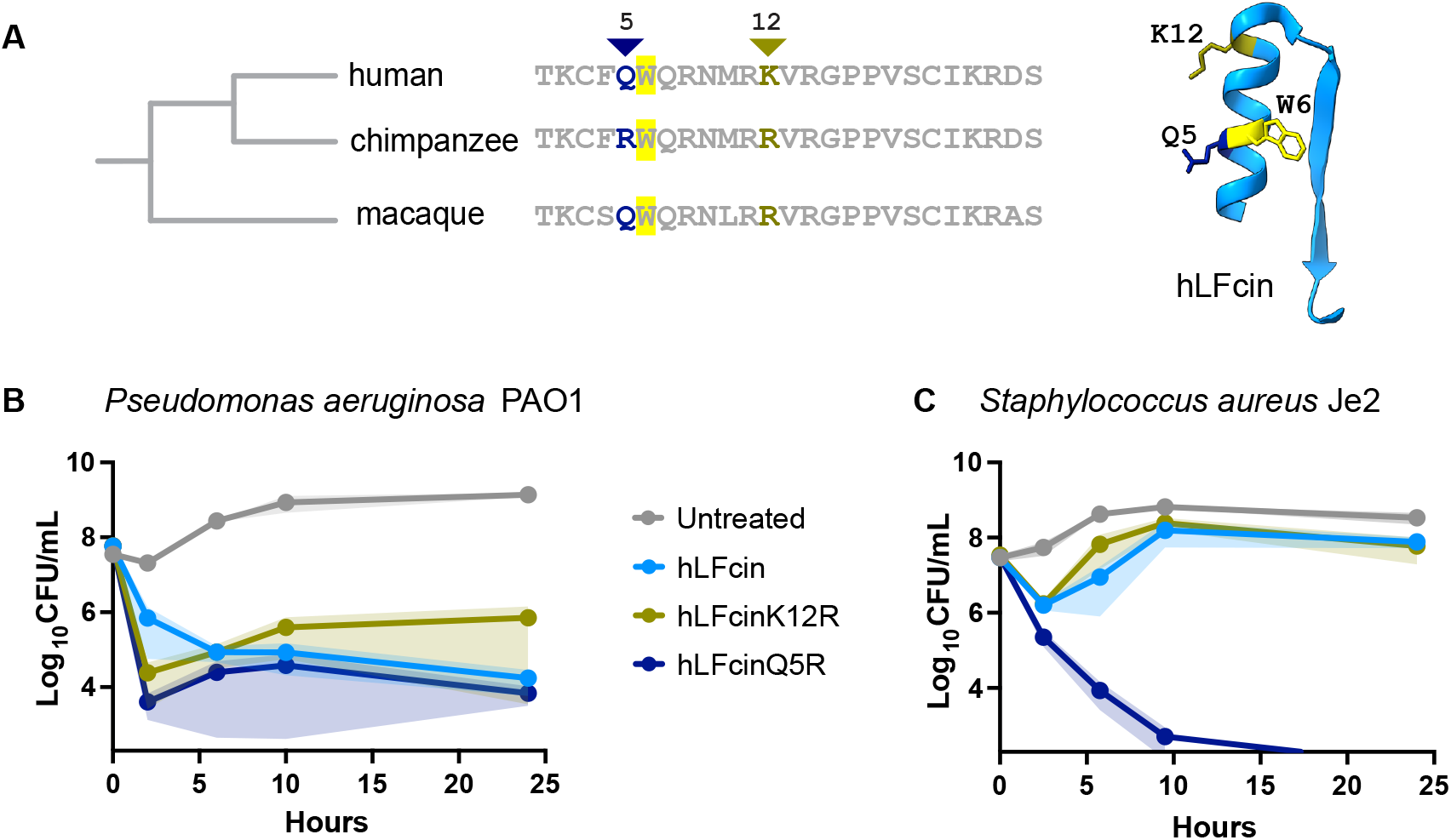
Natural selection enhanced lactoferricin antimicrobial potency in great apes. (A) Alignment of lactoferricin orthologs from humans, chimpanzees, and rhesus macaques. Two sites (positions 5 and 12) exhibit elevated dN/dS across primates. (B) Antimicrobial activity of hLFcin and mutant peptides against *Pseudomonas aeruginosa* PAO1 at 100 μg/mL. (C) Antimicrobial activity of hLFcin and mutant peptides against *Staphylococcus aureus* strain JE2 at 800 μg/mL. Data represent mean and standard error based on three biological replicates.

To assess the functional consequences of natural selection at sites 5 and 12, we generated two point mutants in human lactoferricin: hLFcinK12R and hLFcinQ5R. hLFcinK12R exhibited antimicrobial activity comparable to wild type hLFcin against both *P. aeruginosa* and *S. aureus*, indicating that the K12R substitution does not substantially alter activity against these pathogens (Fig. 5B, C). In contrast, the hLFcinQ5R mutant displayed significant enhancement in antimicrobial potency against *S. aureus* relative to hLFcin (Fig. 5B, C). Beyond the addition of a single positive charge, this effect may stem from the positioning of R5 adjacent to the aromatic hydrophobic residue tryptophan (W6). We noted a similar pattern in AncLFcin2, which exhibited greater potency against *S. aureus* than hLFcin and contains a lysine (K5) adjacent to phenylalanine, another aromatic hydrophobic residue (Fig. 1C, 2C-D, 3D-F). Notably, the most potent variant, bLFcin, contains both combinations, which may contribute to its heightened antimicrobial activity. Taken together, these findings indicate that antimicrobial function of the lactoferricin domain has been repeatedly modulated by the forces of natural selection, leading to enhanced activity across mammalian lineages.

## Discussion

AMPs have long been appreciated as deeply conserved effectors of innate immunity. Recent large-scale analyses further suggest that AMP-like domains are far more widespread than previously appreciated ^10–13^. Computational studies have detected AMP-like domains embedded within diverse proteins not previously associated with direct antimicrobial function, including interferon gamma, IL-13, histone H1, gasdermin E, and dynactin, among others ^10–13^. The existence of such cryptic antimicrobial functions suggests a previously unappreciated layer of host defense, though the evolutionary origins of such embedded AMPs remain largely uncharacterized. Leveraging mammalian lactoferrin and its embedded AMP lactoferricin as a model, we observed that even the earliest lactoferricin ancestor possessed the ability to permeabilize bacterial membranes and alter membrane potential, a property that was further enhanced in later ancestors (Fig. 4). This increased activity was accompanied by a marked perturbation of bacterial cells, likely due to leakage of cytoplasmic content ^49–51^. Membrane permeabilization by AMPs is often reversible depending on the severity of damage ^50,51^. Hence, it is reasonable to speculate that early lactoferricin domains induced perturbations that bacteria could repair, allowing eventual recovery. In contrast, more potent lactoferricin variants such as bLFcin induced irreversible damage at much lower concentrations, leading to complete lysis or severe morphological damage that prevented recovery (Fig. 4). These findings demonstrate that membrane permeabilization and hyperpolarization were early antimicrobial traits of lactoferricin, and that bactericidal activity intensified over time. Assessing the activity of additional lactoferricin ancestors or mutants could aid in further resolving the molecular basis of this activity in future work.

We noted that some ancestral lactoferricins exerted higher antimicrobial activity than their extant orthologs. For example, AncLFcin2 and AncLFcin3 were consistently more bactericidal against *S. aureus* than human lactoferricin (Fig. 2, 3). Thus, the evolution of antimicrobial function in lactoferrin does not reflect a simple linear increase in potency but has rather fluctuated along different lineages depending on the target microbe. We considered whether a reduction in lactoferricin antimicrobial activity could indicate a trade-off to minimize toxicity against host cells. However, treatment of bovine erythrocytes revealed no significant hemolysis in the presence of any lactoferricin peptides, inconsistent with this hypothesis (Fig. S6). An alternative explanation is that some bacteria may have evolved host-specific resistance to these AMPs. *S. aureus*, for example, is human associated with many of its virulence factors and toxins exhibiting narrow tropism for humans ^52–54^. Alternatively, trade-offs with antimicrobial activity could arise due to other factors such as peptide stability, ease of protease digestion, or iron binding ability of the N-lobe. In addition, this study was performed in a low-nutrient laboratory media which do not fully recapitulate the host environment. Future studies could measure the activity of lactoferricin variants in alternative host-mimicking conditions or *in vivo*, which may reveal as-yet unknown impacts on antimicrobial function. Together these results underscore how ancestral sequence reconstruction can uncover unique variants of extant AMPs that could serve as safe and effective pathogen treatment options, particularly in the era of growing antibiotic resistance ^5,55^.

AMPs have frequently been identified among the most rapidly evolving genes in host genomes, likely reflecting repeated adaptation to counteract diverse microbes ^56,57^. In contrast, far less is known about how AMP domains embedded within larger proteins emerge and evolve in response to selection. We previously identified two sites exhibiting signatures of positive selection in the lactoferricin domain among primates ^33^. Notably, one of these substitutions (Q5R) was sufficient to significantly enhance the antimicrobial activity of human lactoferricin, particularly against *S. aureus* (Fig. 5). This observation suggests the importance of this region for lactoferricin’s activity. Notably, the adjacent sixth residue is invariably an aromatic hydrophobic amino acid, either phenylalanine (F) or tryptophan (W), across all lactoferricin sequences (Fig. 1C). AncLFcin2, which is effective against all pathogens tested, also possesses cationic and aromatic hydrophobic residues at these sites (K5F6). The most potent variant tested, bovine lactoferricin, contains three such pairs of cationic and aromatic residues (F1K2, R5W6, and W7R8). Although hTF and AncTF both contain a F6R7 amino acid pair, the effect of the positively charged arginine is likely diminished by the neighboring aspartate (D8). Given that cationic nature and hydrophobicity are key features of many AMPs, the juxtaposition of a cationic residue with an aromatic hydrophobic residue within lactoferrin may reflect positive epistasis between these sites. Indeed, epistasis between nearby R and W residues has been reported previously in antiviral proteins ^58^. It is well established that cationic amino acids mediate electrostatic interactions with the negatively charged bacterial envelopes, whereas hydrophobic residues insert into membranes and contribute to disruption of the lipid bilayer ^59^. Future work could determine how selection and epistasis together contribute to antimicrobial potency in these and other AMPs.

It is notable that lactoferrin has been reported to contain not just one, but multiple embedded antimicrobial domains. For example, the lactoferrin N-lobe contains a 17-amino acid region termed lactoferrampin that possesses antimicrobial activity against both bacteria and fungi ^60^. In addition, ovotransferrin, the ortholog of mammalian transferrin in birds, has also been reported to contain an antimicrobial fragment within the N-lobe in a region distinct from both lactoferricin and lactoferrampin ^61,62^. These observations suggest that embedded AMPs may have arisen multiple times within lactoferrin or transferrin homologs during the course animal evolution. This protein family could thus provide an informative system to further investigate the origins and evolution of antimicrobial functions.

In addition to providing protection against invasive pathogens, the expression of AMPs at barrier tissues positions them to broadly shape host-associated microbiotas. Work in *Drosophila* has demonstrated how natural selection and polymorphism in AMP genes can impact diverse resident microbes ^56,63,64^, as well as confer defense against microbes within specific ecological niches ^65^. Given that lactoferrin is among the most abundant secreted proteins in many mammalian barrier tissues, genetic variation may similarly shape the diversity and composition of host-associated microbial communities. By retracing the origin and evolution of antimicrobial activity in mammalian lactoferrin, our study illustrates how new protein functions can emerge to promote host defense against major microbial pathogens.

## Materials and Methods

### Ancestral sequence reconstruction

Vertebrate transferrin and mammalian lactoferrin homologs were retrieved from the NCBI and UniProt databases for ancestral sequence reconstruction. Multiple sequence alignments and maximum likelihood phylogenetic trees were generated using the Topiary pipeline ^37^. Alignments were manually scanned to remove insertions and deletions. Using Topiary, ancestral sequences were inferred at each node, with posterior probabilities calculated for each site and node to assess reconstruction confidence.

### Peptide synthesis

All peptides were commercially synthesized by GenScript. Lyophilized peptides were resuspended in phosphate-buffered saline (PBS, pH 7.0) to a concentration of 10 mg/mL and stored at –80°C.

### Structural prediction of ancestral sequences

Protein structure predictions for the reconstructed ancestral sequences were performed using ColabFold, a streamlined implementation of AlphaFold2^66^. Prior to structure prediction, the N-terminal region corresponding to the human lactoferrin signal peptide (hLF1–19) was removed from all ancestral sequences. Predicted protein structures were visualized and analyzed using UCSF ChimeraX (version 1.8, released 2024-06-10)^67^. The theoretical isoelectric point (pI) of each peptide was calculated using the ExPASy Compute pI/Mw tool^68^.

### Bacterial growth assays

Overnight cultures of all bacterial strains were prepared by inoculating a single colony into low-nutrient 5% tryptic soy broth (TSB, v/v) to model the low-nutrient conditions of the host environment (*5*) and incubated at 250 rpm overnight, unless otherwise specified. The following day, cultures were diluted to an optical density at 600 nm (OD_600_) of 0.02 in fresh 5% TSB. For growth assays, 150 μL of the diluted culture was added to each well of a 96-well microtiter plate, followed by the addition of peptides at the desired concentrations. Plates were incubated for 24 hours with continuous shaking in a Synergy H1 microplate reader (BioTek), and bacterial growth was monitored by measuring OD_600_ at 10-minute intervals. Bacterial growth was then normalized by subtracting the optical densities of similar volume of media with or without the peptides of particular concentrations from those of bacterial cultures. Area under the curve (AUC) was calculated and normalized on GraphPad Prism software. For normalization of the AUCs, 100% was defined as the average measurements from the bacterial growth without any peptide added.

### Bacterial survival assay

Overnight cultures were diluted to an OD_600_ of 0.02 in fresh medium, and antimicrobial peptides were added at the indicated concentrations in 1.5 mL microcentrifuge tubes. To assess bacterial viability over time, samples were taken at defined time points during the treatment period. At each time point, cultures were serially diluted 10-fold (up to 10^−7^) and plated on tryptic soy agar (TSA) to enumerate colony-forming units (CFUs). Plates were incubated at 37°C overnight, and CFUs were counted the following day to evaluate the time-course of bacterial survival.

### Membrane permeabilization measurements

Overnight cultures of *Pseudomonas aeruginosa* PAO1 were subcultured by diluting to an OD_600_ of 0.3 in fresh 5% TSB. Aliquots of 100 μL were transferred into 1.5 mL microcentrifuge tubes and treated with the peptides for 30 minutes. Cells were pelleted by centrifugation at 5,000 × g for 1 minute. Pellets were resuspended in 300 μL of 20 μM propidium iodide (Sigma-Aldrich P4170-10MG) and incubated in the dark for 15 minutes to prevent photobleaching. Excess dye was removed by centrifugation at 5,000 × g for 1 minute, the supernatant was discarded, and pellets were resuspended in 300 μL of phosphate-buffered saline (PBS, pH 7.0). For fluorescence measurement, 100 μL aliquots from each sample were transferred in triplicate into black-walled, clear-bottom 96-well plates (without lids). Fluorescence intensity was measured from the bottom of the plate using a microplate reader with excitation at 535 nm and emission at 617 nm.

### Membrane potential measurements

The membrane depolarizing activity of peptides was assessed using the voltage-sensitive dye DiSC_3_(5) (MedChem Express #HY-D0085-25mg) as previously described^48^. *Pseudomonas aeruginosa* PAO1 overnight cultures were diluted in fresh 5% TSB to OD_600_ of 0.3. DiSC_3_(5)^48^, prepared as a stock solution in DMSO, was added to the bacterial suspension at a final concentration of 5 μM. Following a brief equilibration period to allow dye accumulation across polarized membranes, peptides were added at a final concentration of 100 μg/mL in a black, clear-bottom 96-well plate. Fluorescence was monitored kinetically over a 1-hour period at 37 °C using a Synergy H1 microplate reader (BioTek), with excitation and emission wavelengths set to 610 nm and 660 nm, respectively. A decrease in fluorescence intensity was interpreted as membrane hyperpolarization, reflecting increased accumulation of the cationic, lipophilic dye within the bacterial cytoplasmic membrane, where self-quenching reduces the overall fluorescence signal.

### Nile red staining, microscopy and cell size measurements

The Nile red staining protocol was adapted and modified from Dombach *et al*., 2021^69^. Overnight bacterial cultures were diluted in fresh 5% tryptic soy broth (TSB, v/v) to an optical density at 600 nm (OD_600_) of 0.3. From this, 100 μL aliquots were transferred to 1.5 mL microcentrifuge tubes for peptide treatment. Samples were incubated at 37 °C for 40 minutes with shaking at 250 rpm. Following peptide treatment, Nile red (Sigma-Aldrich 72485-100MG) was added to each tube at a final concentration of 10 μg/mL, followed by a 5-minute incubation at 37 °C. Cells were then fixed by adding paraformaldehyde to a final concentration of 4%, and incubated at room temperature for 10 minutes. After fixation, cells were pelleted by centrifugation at 10,000 × g for 1 minute and resuspended in 10 μL of PBS, pH 7.0. For microscopy, 7 μL of the resuspended cells was applied to a coverslip, overlaid with 0.5% agarose, and gently compressed with a glass slide to immobilize the sample. Imaging was performed using a SoRA spinning disk confocal microscope (Nikon CSU-W1) equipped with a 60X water-immersion objective. Bacterial cell size was measured using the Fiji package of ImageJ software^70^.

### Hemolysis assay

The hemolysis assay was adapted from Greco *et al*., 2020^71^. Bovine red blood cells (RBCs) were isolated by centrifuging 1 mL of whole blood at 3000 × *g* for 2 min. The RBC pellet was washed five times with phosphate-buffered saline (PBS) by resuspension and centrifugation until the supernatant was clear, then diluted 1:10 in PBS. In a 96-well plate, 100 µL of the diluted RBC suspension was mixed with each peptide solution (final concentration: 1 mg/mL) and incubated at 37 °C for 1 h. Absorbance was measured at 405 nm using a microplate reader, and the percentage of hemolysis was calculated relative to a positive control (1% Triton X-100).

## Acknowledgements

We thank Mike Harms and Jose Sanchez-Borbon for assistance with ancestral sequence reconstruction. We are thankful to the members of Barber lab for their feedback and support, as well as Karen Guillemin, Jarrod Smith, and Melanie Spero for helpful insights and discussion. We thank the Genomics and Cell Characterization Core Facility (GC3F) at UO and Adam Fries for assistance with microscopy.

