## Supplemental figures for "Stepwise origin and evolution of a cryptic antimicrobial peptide in mammalian lactoferrin"

**This PDF file includes:**

Supporting text
Figures S1 to S6
Legends for Datasets S1 to S2

**Other supporting materials for this manuscript include the following:**

Datasets S1 to S2

**Supplementary figures**

A

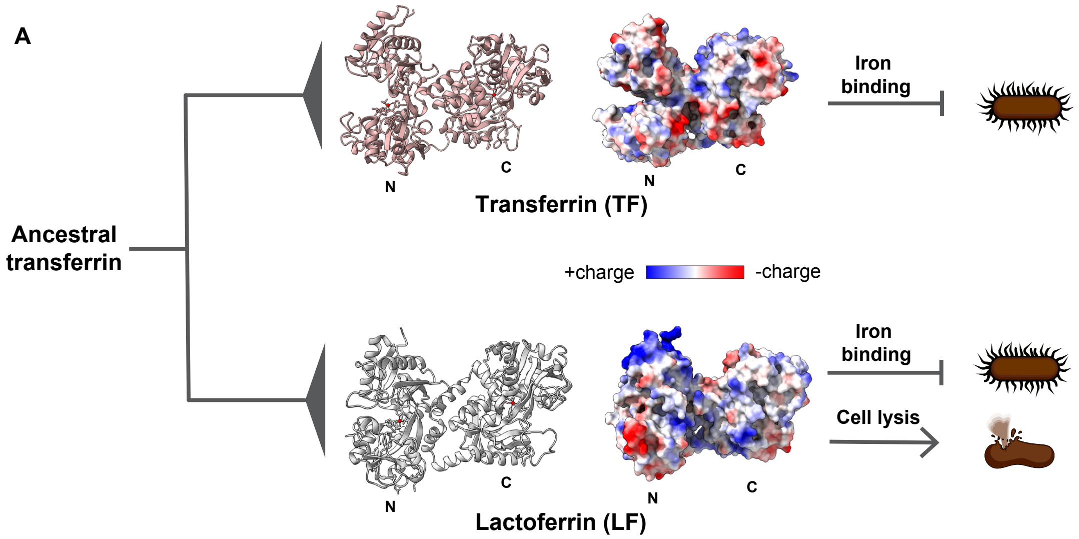


B

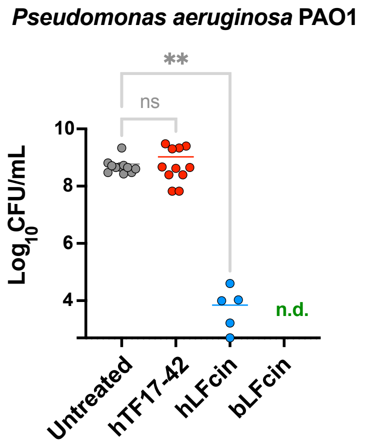


**Fig. S1. Diversity and evolution of lactoferricin sequences.** (**A**) The extant lactoferrin (PDB: 1lfg) and serum transferrin (PDB: 3qyt) genes arose via duplication of transferrin in the ancestor of placental mammals. Surface electrostatic potential is shown. While transferrin maintains key roles in iron binding and transport, lactoferrin has acquired new functions including direct bactericidal activity. (**B**) Antimicrobial activity of extant human and bovine lactoferricin (hLFcin and bLFcin, respectively) against *Pseudomonas aeruginosa* PAO1 compared with the equivalent region in human transferrin (hTF17-42).

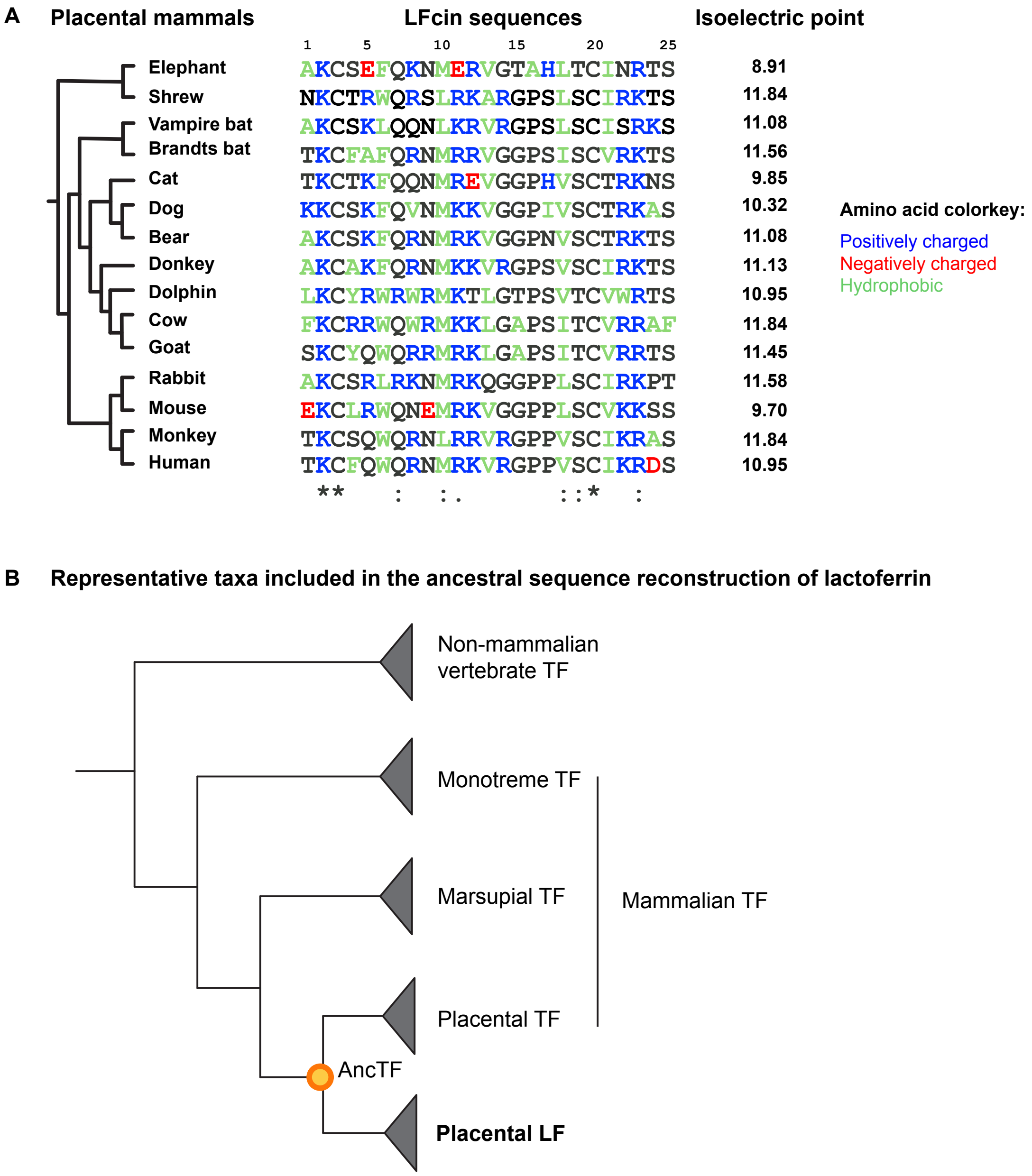


**Fig. S2. Ancestral sequence reconstruction (ASR) of vertebrate transferrin family proteins.** (**A**) Amino acid alignment of the 25-amino acid stretch of lactoferricin region across placental mammals. Blue, red and green letters denote positively charged, negatively charged, and hydrophobic aromatic residues, respectively. (**B**) A total of 376 sequences of transferrin, lactoferrin, and melanotransferrin from diverse vertebrate species were used to reconstruct the ancestral sequences.

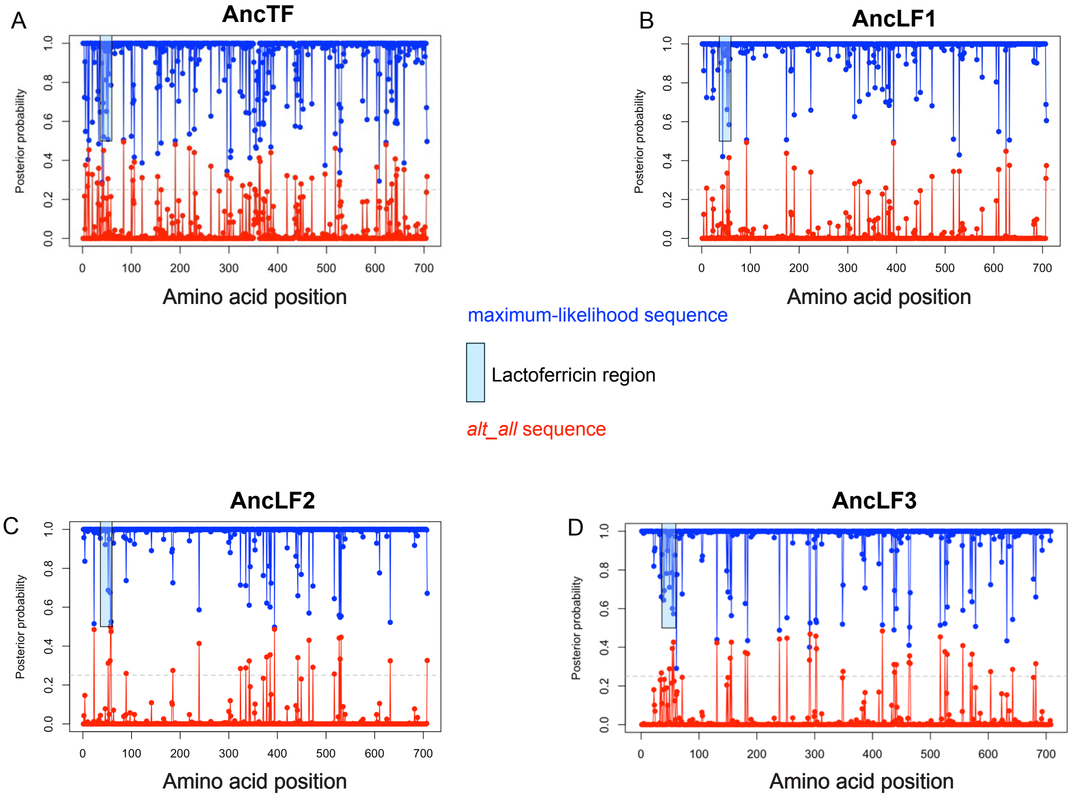


E

| | |
|---|---|
| AncTF17-42 | SKCSSFRDNMKKADGPSVTCVKRTS |
| Altall_AncTF17-42 | SKCSKFRENMKKADGPPVTCVKRTS |
| AncLFcin1 | AKCSKFQQNMKKVGGPSVTCIKRTS |
| Altall_AncLFcin1 | AKCSKFQENMKKVGGPPVTCVKRTS |
| AncLFcin2 | AKCSKFQRNMKKVGGPSVSCIKKTS |
| Altall_AncLFcin2 | AKCSKFQRNMKKVGGPPVSCVRRTS |
| AncLFcin3 | SKCSQWQRRMKKVGGPSVTCVRKTS |
| Altall_AncLFcin3 | SKCSQWQRRMKKVRGPSVSCIRKTS |

**Fig. S3. Reconstructed ancestral sequences and alternate variants.** Posterior probabilities of each site are shown for (**A**) AncTF, (**B**) AncLF1, (**C**) AncLF2 and (**D**) AncLF3. Blue indicates the most probable amino acid at each position, while red represents the *alt_all* sequence containing the second-most probable amino acid at each site. The lactoferricin regions are highlighted with cyan rectangles in the full-length sequences. (**E**) Sequences of the lactoferricin regions in the reconstructed ancestors. Yellow highlight denotes residues that differ between the most probable (blue) and *alt_all* (red) sequences.

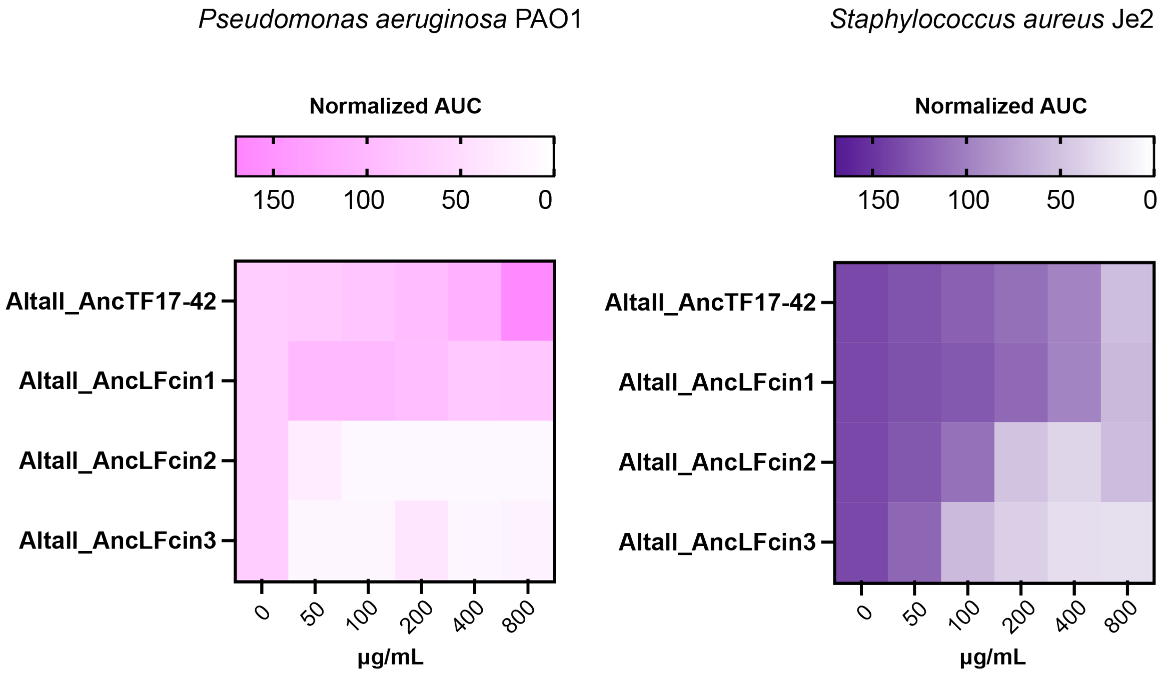


**Fig. S4. Antimicrobial activities of alternate ancestral peptides.** Heatmaps show area under the curve (AUC) values normalized against the untreated control from bacterial growth curves in the presence of the indicated peptides at specified concentrations. Gram-negative strains are shown in pink, Gram-positive strains in purple.

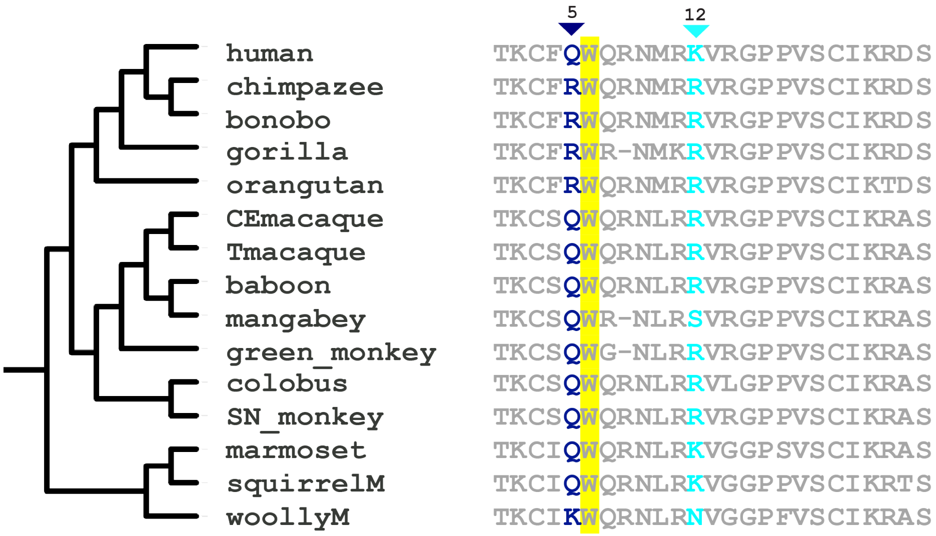


**Fig. S5. Evolution of lactoferricin across simian primates.** Lactoferricin positions 5 and 12 exhibited elevated dN/dS consistent with repeated positive selection. In Great Apes, glutamine (Q) at position 5 is replaced by arginine (R), adjacent to a tryptophan (W). At position 12, residues vary between arginine (R) and lysine (K) across primate species.

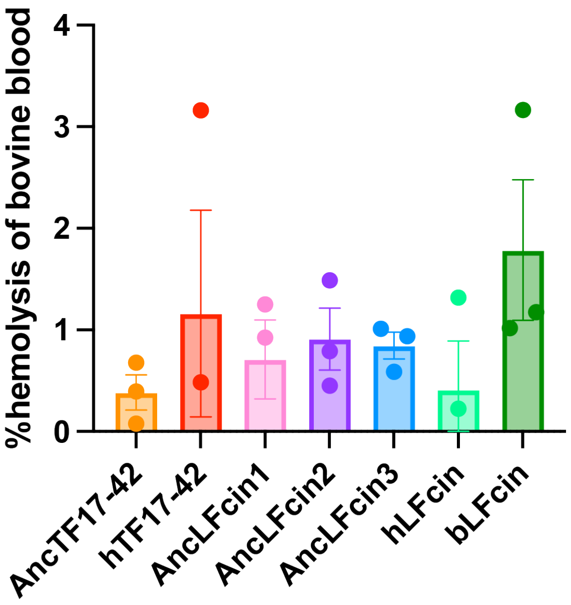


**Fig. S6. Hemolysis of ancestral and extant peptides.** Bovine red blood cells (RBCs) were treated with peptides at 1 mg/mL. Complete lysis (100% hemolysis) was achieved with 1% Triton X-100 as a positive control. Hemolysis by the peptides is shown as a percentage relative to this control. Data represent three biological replicates with the standard error.

**Dataset S1 (separate file). Newick file of the ancestral sequence reconstruction tree.** The tree contains 376 vertebrate sequences of transferrin (TF), lactoferrin (LTF) and melanotransferrin (MLTF) used to infer the ancestral sequences.

**Dataset S2 (separate file). Reconstructed ancestral sequences used for analysis.** This file contains the full-length amino acid sequences of AncTF, AncLF1, AncLF2, and AncLF3, including both their most likely sequences and the *alt_all* variants, in which each position is assigned the second most likely amino acid. The average posterior probability (PP), number of ambiguous sites (num_ambig), and number of ambiguous gaps (num_ambig_gaps) for each sequence are provided in the sequence headers. The magenta-highlighted region (first 19 amino acids) represents the predicted signal peptide, which was removed prior to structural visualization. The cyan-highlighted region corresponds to the human lactoferricin-equivalent segment of 25 amino acids, which was used for experimental purposes to test the evolutionary development of antimicrobial activity within the lactoferricin domain.
